# Unravelling the binding mode of a methamphetamine aptamer: a spectroscopic and calorimetric investigation

**DOI:** 10.1101/2021.08.13.456068

**Authors:** Clement Sester, Jordan AJ McCone, Ian Vorster, Joanne E Harvey, Justin M Hodgkiss

## Abstract

Nucleic acid aptamers are bio-molecular recognition agents that bind to their targets with high specificity and affinity, and hold promise in a range of biosensor and therapeutic applications. In the case of small molecule targets, their small size and limited number of functional groups constitute challenges for their detection by aptamer-based biosensors because bio-recognition events may both be weak and produce poorly transduced signals. The binding affinity is principally used to characterize aptamer-ligand interactions; however a structural understanding of bio-recognition is arguably more valuable in order to design a strong response in biosensor applications. Using a combination of nuclear magnetic resonance, circular dichroism, and isothermal titration calorimetry, we propose a binding model for a new methamphetamine aptamer and determine the main interactions driving complex formation. These measurements reveal only modest structural changes to the aptamer upon binding and are consistent with a conformational selection binding model. The aptamer-methamphetamine complex formation was observed to be entropically driven, apparently involving hydrophobic and electrostatic interactions. Taken together, our results establish a means of elucidating small molecule-aptamer binding interactions, which may be decisive in the development of aptasensors and therapeutics, and may contribute to a deeper understanding of interactions driving aptamer selection.

## INTRODUCTION

Nucleic acid aptamers are functional bio-molecular recognition agents that bind to their targets with high specificity and affinity (1). They mostly originate from *in vitro* selection experiments involving the systematic evolution of ligands by exponential enrichment (SELEX) method which, starting from random sequence libraries, optimizes the nucleic acid sequences for high-affinity binding to the presented target ligands (2,3). Aptamers have proven to be highly stable, readily adaptable to chemical modification, and exhibit reversible binding. As a consequence, aptamer-based biosensors (aptasensors) are promising replacements for antibody-based biosensors in many applications (4,5).

An understanding of the aptamer-ligand binding domain and mechanism is valuable for sensing applications. With this knowledge, the design of the specific binding pocket by truncation or addition of nucleotides, or hybridization with a complementary nucleotide sequence can result in enhancement of the specificity and the signal transduction, and thus the sensitivity of the sensor (6–9). Moreover, many sensing schemes rely on substantial target-induced structural changes to the aptamer. In this case, careful placement of a label can induce an electrochemical or fluorescent signal (10,11). On the other hand, even an aptamer-ligand complex with a strong binding constant may not make a good sensor if a lack of structural change prevents strong signal transduction. To a large extent, the degree of structural change upon ligand binding is expected on thermodynamic grounds to be negatively correlated with the binding constant. In any case, the most appropriate transduction system for a given aptamer-ligand pair can be selected according to the thermodynamic and structural properties of its complex (10,12).

The binding dynamics of aptamer-ligand systems can be described by three binding models, derived from the framework of protein-ligand interactions that describe folding tunnels and free energy basins (Figure 1) (6,13–15).

**Figure 1.**
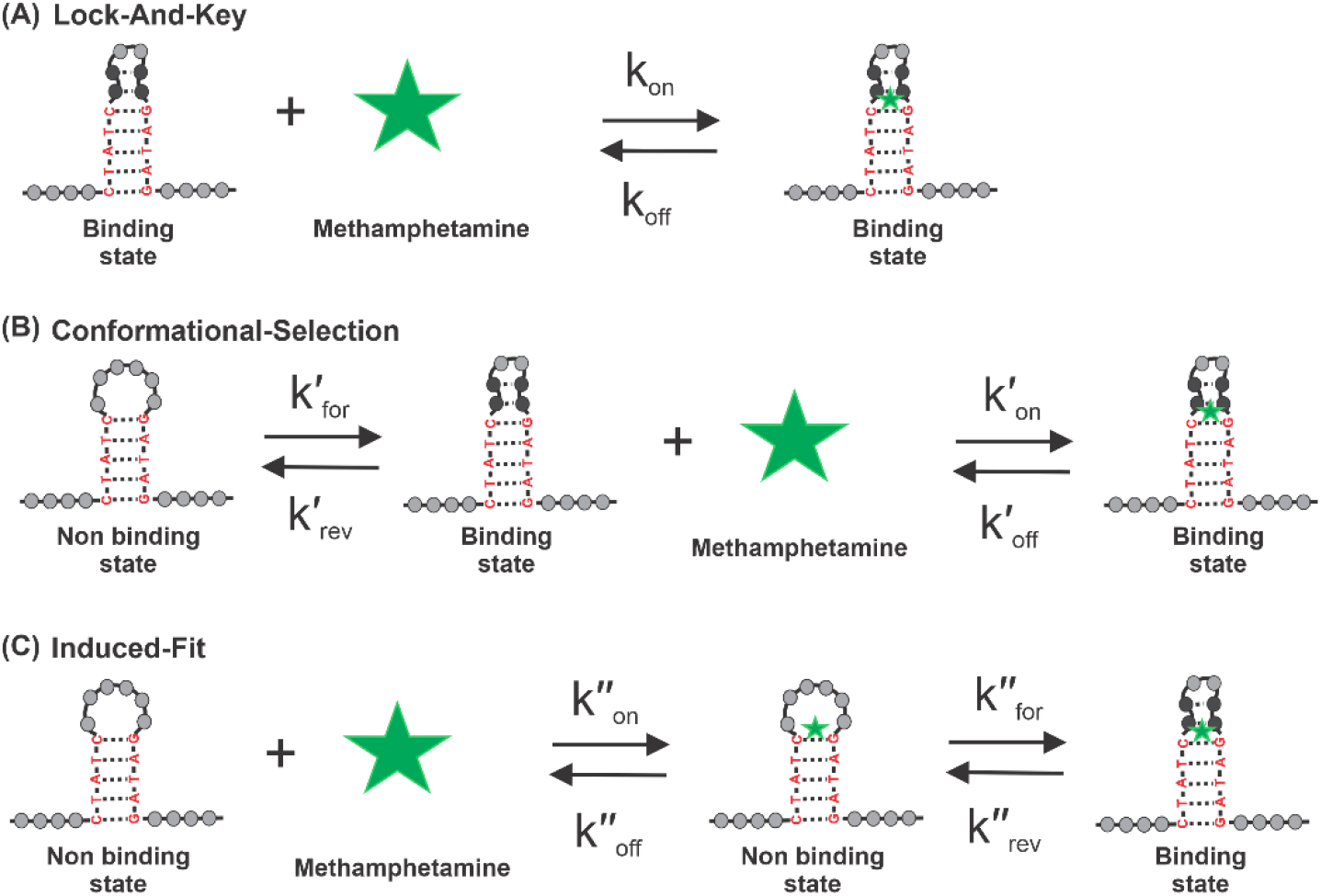
Binding models of bioreceptors in the context of aptamer-ligand binding. (A) Lock and Key (LAK), (B) Conformational Selection (CS) and (C) Induced Fit binding models. Representation relates to the interaction of the Aptamer-2-40mer with methamphetamine. Binding constant equations are found in Equation S1.

The lock-and-key (LAK) binding model (Figure1A) simplifies the binding interaction as between a rigid bio-receptor and ligand (16). Consequently, their binding interfaces are assumed to be perfectly matched. Accounting for this assumption in the case of aptamer-ligand interactions, the specific nucleotide strand is modeled as though it undergoes no structural change upon ligand addition. π-Stacking, van der Waals interactions, and hydrogen bonding between the aptamer and the ligand stabilize the complex formed (17).

The induced-fit (IF) binding model (Figure 1B) considers the aptamer to have a flexible binding site able to adopt the optimal binding conformation (or structural ‘switch’) for a ligand (18). Therefore, this model does not require the initial binding interfaces between the aptamer and ligand to match. The structural switch causes the formation of Watson-Crick or non-Watson-Crick base pairings in the aptamer structure, which enables the arrangement of a specific binding interface for the ligand. In addition to the aptamer-target interactions considered in the LAK binding model, the formation of new intra-aptamer hydrogen bonds is possible (13).

The conformational-selection (CS) binding model (Figure 1C) acknowledges the inherent dynamic behavior of the aptamer in solution. Numerous conformational states coexist in equilibrium with different population distributions; the ligand can bind selectively to one of the conformational states, ultimately shifting the equilibrium towards this state (19).

In reality, aptamers are known to retain structural plasticity, and bind to their targets by changing their structure in accordance with the target structures (17). In seeking to develop aptamers for small-molecules sensing applications, a significant and measurable biorecognition event is required. In this context, it is important to know whether the aptamer-ligand interaction is necessarily accompanied by a structural change. If so, the timing of the conformational change in relation to the binding event must be determined. Furthermore, small-molecule ligands, by virtue of negligible size and few functional groups, may not induce a significant structural change. Finally, the focus on the dissociation constant, *KD*, for sensing experiments remains valuable, but the crucial requirement for detection by sensors is the measurable transduction of the bio-recognition event. Without specific and significant structural change, a reliable signal may not be detected.

Here, we propose an aptamer characterization workflow using spectroscopic and calorimetric methods coupled with *in silico* predictions to get a more complete understanding of the interaction between an aptamer and its small molecule ligand. This workflow starts with the determination of the overall complex tertiary structure parameters and concludes with an in-depth analysis of the specific region and binding interactions between the aptamer and the ligand. Altogether, this information allows us to propose a detailed binding mechanism for the interaction between a small molecule and a SELEX-generated aptamer.

We focus on an aptamer selected to bind methamphetamine (Meth) – a controlled substance that produces a psychological and physical stimulant effect and can lead to dependency. The simple structure of Meth makes it relatively easy to produce illegally, which can also lead to significant issues of contamination in properties used for clandestine manufacturing (20).

The combination of nuclear magnetic resonance, circular dichroism, and isothermal titration calorimetry measurements allows us to propose a binding model for the new Meth aptamer and determine the main interactions during complex formation. Little structural change in the aptamer was detected, and a conformational-selection binding model was proposed. Entropically driven formation of the aptamer-Meth complex, combining hydrophobic and electrostatic interactions, was observed. Molecular docking simulations support the findings made with NMR and ITC.

This approach should be applicable to any aptamer selected for binding a small molecule, enabling a better structural understanding of the complex formed. In the context of aptasensor development, the structural information collected will help in the design and choice of the most suitable transduction platform.

## MATERIALS AND METHODS

### Materials

AuramerBio (Wellington, New Zealand) selected the 75 nucleotides long Meth aptamer family via “Affinity Matrix Selex” (21). Synthetic oligonucleotides were obtained from AlphaDNA (Montreal, Canada). DNA strands were purified by SDS-PAGE. The binding washing buffer (BWB) was tris(hydroxymethyl)aminomethane hydrochloride buffer (2 mM Tris-HCl, 10 mM NaCl, 0.5 mM KCl, 0.2 mM MgCl_2_ and 0.1 mM CaCl_2_; pH 7.5). Sodium 3-(trimethylsilyl)-1-propanesulfonate (DSS) was used as an internal standard for NMR experiments (22). Before any experimentation, the aptamer was denatured at 95 °C for 5 min followed by cooling in an ice bath for 10 min.

### SYBR green assay I

Each aptamer was mixed in BWB solution with the fluorescent dye SG then titration with Meth was conducted for each of them; fluorescence signal is monitored during the Meth titration. 10 μl of SG (7x) and 10 μl of aptamer (10 μM) were mixed. A range of Meth concentrations varying from 0 to 1 μM was prepared in BWB and added directly to the SG-aptamer solution to deliver a final volume of 1000 μL. Fluorescence emission spectra were recorded from 500 to 650 nm using an excitation wavelength of 497 nm. The fluorescence emission of SG-stained DNA was measured at 520 nm.

### CD analysis

BWB solutions containing each aptamer with and without methamphetamine were analyzed with a CD instrument. BWB solutions of each aptamer (1.5 μM) were prepared with and without Meth (3 μM). CD analyses were performed on a Chirascan Spectrometer in a 1 mm path length quartz cuvette. The spectra were recorded from 380 to 180 nm at 20 °C at a rate of 200 nm/min and corrected by subtraction of the background scan of the buffer.

### ^1^H NMR and Nuclear Overhauser enhancement (NOESY) experiments

The aptamer sample was analyzed by ^1^H NMR and 2D NOESY. DNA samples were dissolved in 527 μL of BWB containing 10 % D_2_O. The final concentration of DNA was 1 mM. An internal standard DSS was added to the sample and its final concentration was 1 mM. For spectra in the presence of Meth, the Meth concentration was fixed to 1 mM. NMR experiments were performed on a JEOL 600 MHz Nuclear Magnetic Resonance Spectrometer type JNM-ECZ600R with data recorded at 20 °C. 1D proton NMR spectra were acquired with 3500 data points using 1280 scans due to low sample concentration. NOESY spectra were obtained using a mixing time of 200 ms (23). Solvent suppression was achieved by the “Watergate sequence” (24). Acquired data were processed and analyzed using MestReNova software (25).

### Isothermal titration calorimetry assay

Calorimetric experiments were conducted with the aptamer and a stem mutated version. Isothermal titration calorimetry (ITC) assays were performed on an Affinity ITC system (TA Instruments Inc., New Castle, DE, USA). 1 μL injections of a 2.5 mM Meth solution in BWB were added by a computer-controlled microsyringe at an interval of 240 s into the DNA aptamer solution (50 μM in BWB, cell volume 200 μL) with 100 rpm stirring at 20 °C. Heats produced by Meth dilution and aptamer dilution were evaluated and subtracted. The experimental data were fitted to a theoretical titration curve using software supplied by TA, with *ΔG* (Gibbs free energy), *K_D_* (dissociation constant), and *n* (number of binding sites per monomer), as adjustable parameters.

### Molecular modeling

Three plausible two-dimensional (2D) structures of Aptamer-2-40mer were generated using Mfold and RNAstructure (26,27). The 2D structure, related to the lowest Gibbs free energy, was then imported to RNAComposer (28) to generate a three-dimensional (3D) model, which was used in subsequent docking studies. The 2D structure of Meth was sketched on Maestro and prepared for docking by generating a low energy 3D conformer using LigPrep (29). The program GOLD (Genetic Optimization for Ligand Docking) (30) was used for all molecular docking analyses. The aptamer binding site was defined as a 100 Å sphere around the coordinates x = 0.3155 y = −26.7188 z = 6.1465. Because GOLD’s GA (genetic algorithm) is stochastic in nature, an exhaustive approach was adopted such that 100 GA runs were set on the input conformation, giving 100 poses for further analysis. Search efficiency was set to 200 % (highest accuracy). The ChemPLP (31) scoring function was used to score and rank poses, with a higher score corresponding to a pose having a relatively better ‘fit’ to the cavity. The resulting 100 poses were clustered according to their respective geometries using the *cluster_conformers.py* script implemented in the Schrödinger 2019-2 package (32). The average linkage method was used for cluster generation and the optimal number of clusters was automatically determined using the Kelley index (33). Finally, clusters were visually compared and merged based on their conformational similarity.

## RESULTS AND DISCUSSION

The 75-nucleotide long methamphetamine (Meth) aptamer family developed via Affinity Matrix SELEX comprised four structures: Aptamer-1, Aptamer-2, Aptamer-3, and Aptamer-4 (Table S1). At pH 7.5, the cationic form of Meth dominates (Figure 2), making the ammonium part of the structure hydrophilic and electron deficient (red shading), while the phenylpropyl moiety is hydrophobic and electron-rich (grey shading). The hypothetical interactions with the DNA strand of the aptamer will be electrostatic attractions with the cationic nitrogen, hydrogen bond formation between the protonated amine and an atom of oxygen or nitrogen in the aptamer, and hydrophobic and π-stacking with the phenyl ring (17,34).

**Figure 2.**
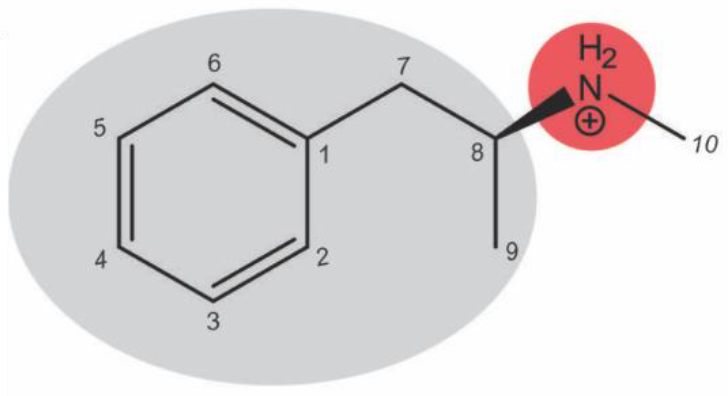
Skeletal formula of protonated Meth, as dominant at pH 7.5.

A fluorescence experiment (SYBR I green [SG] assay) was undertaken to compare the binding affinity of the four aptamers with Meth and to guide the choice between the four aptamers for further structural investigations (Figure S1). The SG molecule emits a fluorescent light by interacting non-covalently with the duplex part of a DNA strand (35). The addition of the target molecule to the SG-aptamer complex causes the release of the SG molecule from the DNA structure induced by the interaction between the target and the DNA strand. As a consequence, a decrease of intensity of the fluorescent light can be monitored, and the interaction event between the aptamer and its target, as well as the resulting *K_D_*, can be deduced (36). Based on the best *K_D_* found with the SG assay (*K_D_* = 244.2 nM) and on the structural information acquired with CD experiments (Figure 3 and Figure S2), Aptamer-2 was chosen for a more thorough structural investigation.

**Figure 3.**
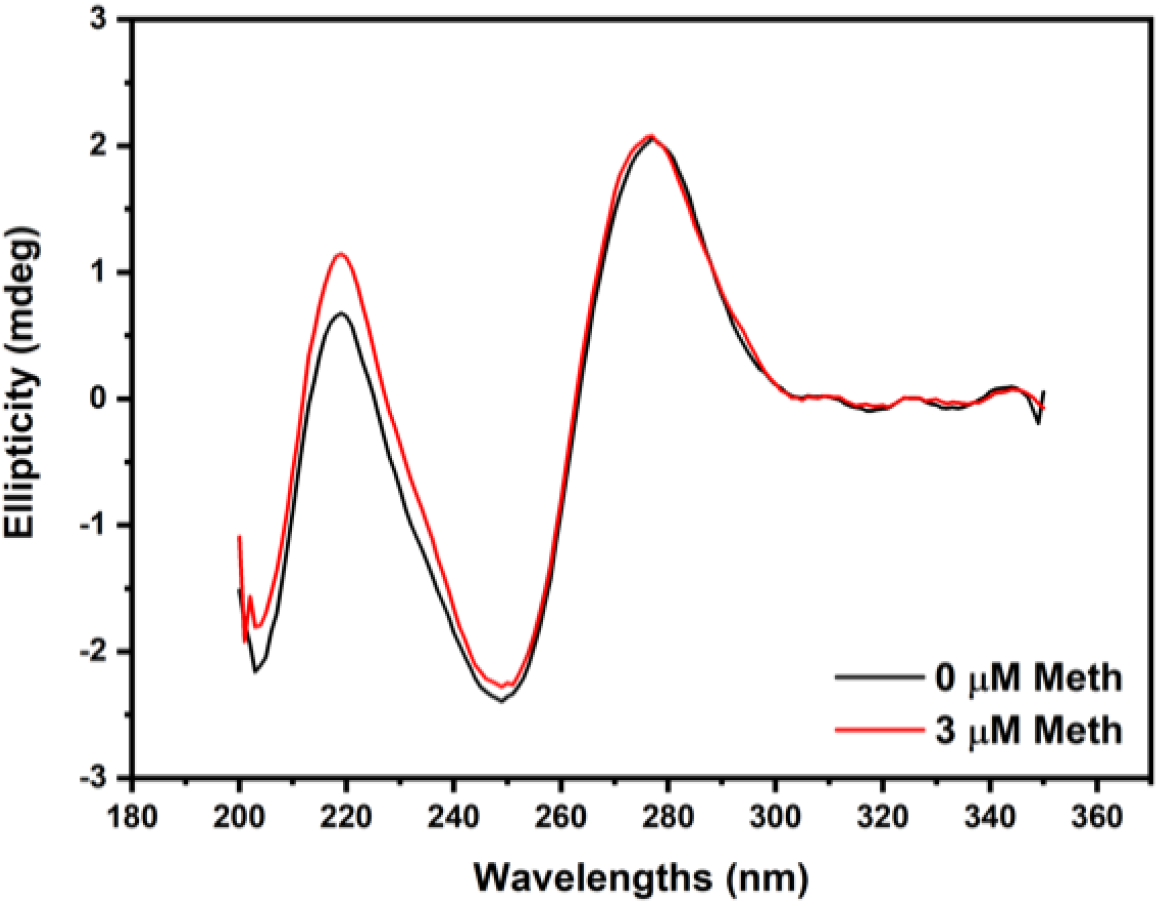
CD experiments for Aptamer-2 upon Meth addition. The positive ellipticity peak at 277 nm and the negative ellipticity peak at 249 nm are characteristic of a B-form structure.

### Circular Dichroism (CD) shows that Aptamer-2, in the presence or absence of Meth, has a B-form structure

The CD spectrum (Figure 3) of Aptamer-2 reveals a positive ellipticity peak at 277 nm and a negative ellipticity peak at 249 nm, characteristic of a B-form structure containing a duplex region (37–39). This is assumed to be the most stable form of the aptamer at the conditions used for the experiment (40,41). Heterogeneous nucleotide DNA sequences typically adopt the B-form structure, in contrast to RNA strands that adopt an A-form structure (which produces a positive band at 260 nm and a negative band at 210 nm), or guanine-rich sequences that will preferentially adopt G-quadruplex structures (producing two positive bands at 215 and 295 nm and a negative band at 260 nm) (42). In our case, the addition of Meth to Aptamer-2 caused no shift or significant changes in peak intensity for the two distinguishable B-form peaks (Figure 3). This indicates that Aptamer-2 retains its B-form conformation upon target addition. Regarding the binding models, this could indicate that the major secondary structural motif is preformed and is not dependent on the presence of the target, meaning the IF binding model could be ruled out.

### Computational predictions of the truncated Meth aptamer, Aptamer-2-40mer, disclose a potential secondary structure

To obtain simplified NMR spectra for analysis, a truncated aptamer variant (reduced from 75 to 40 nucleotides) was used: Aptamer-2-40mer. The non-essential parts (5′ primer and 3′ primer) of the original Aptamer-2 were removed. The primers are necessary to amplify target-bound sequences by PCR during the SELEX process. They do not belong to the randomized part of the aptamer and thus can be removed without significant effect on ligand binding (43). Additionally, they can convolute results by interacting non-specifically with the ligand and so their removal is desirable for our subsequent studies.

For Aptamer-2-40mer, three secondary structures were predicted computationally by Mfold and RNAstructure (26,27). All three structures contained a hairpin fold with duplex region. The form represented in Figure 4 was calculated by Mfold to have the lowest Gibbs free energy at 20 °C and at the specific buffer ionic strength used. Moreover, the structure shown in Figure 4 is consistent with that predicted by Mfold for the original 75mer version of the Aptamer-2. NMR analysis was undertaken to confirm the presence of the hairpin loop structure.

**Figure 4.**
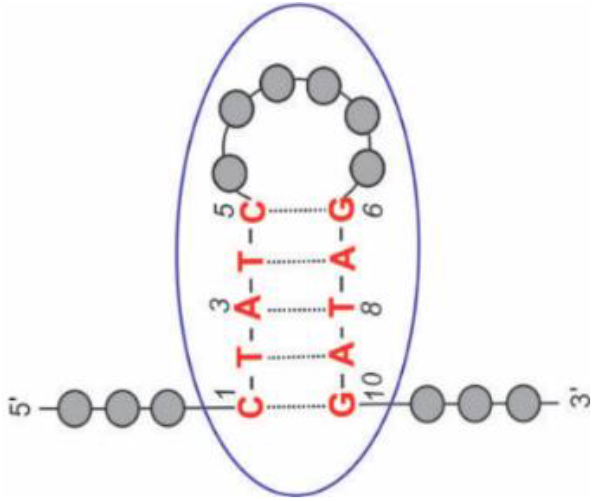
Representation of the Mfold-calculated lowest energy Aptamer-2-40mer structure with the hairpin structure circled.

### NMR experiments with Aptamer-2-40mer partially identify the nucleotides belonging to the stem

The ^1^H NMR spectrum of Aptamer-2-40mer (Figure S3) is typical of a partially double-stranded DNA sequence. Peaks observed in the 9–15 ppm region correspond to imino protons in the thymine and guanine bases (44,45). Due to proton exchange between imino protons and bulk solvent, the only imino protons typically observed are those involved in hydrogen bonding (H-bonding) through nucleotide base pairing, which can include Watson-Crick (Figure 5) or non-Watson-Crick interactions, or otherwise protected by structural folding (44–46). These types of interactions influence the tertiary structure that may contain a solvent-exposed pocket capable of binding proteins and small molecules (47,48). The resonance region of the Watson-Crick imino protons in an ^1^H NMR spectrum is 12–15 ppm and the resonance region of non-Watson-Crick imino protons is 9–12 ppm (44,45).

**Figure 5.**
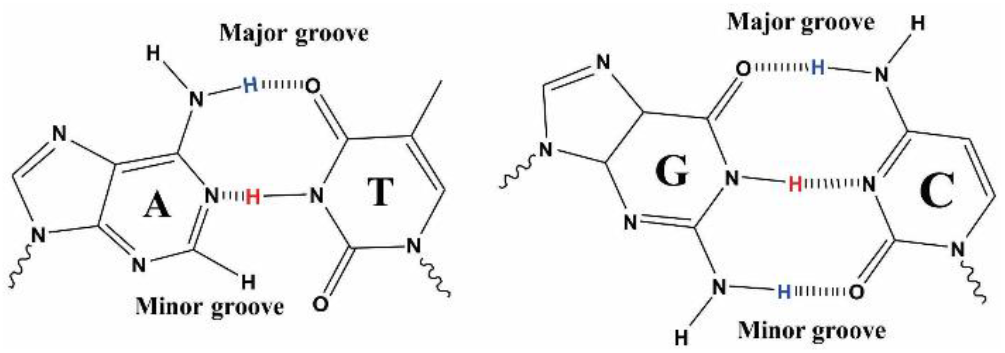
Watson-Crick base pairs A-T and G-C.

Aptamer-2-40mer contains five Watson-Crick base pair interactions (two G-C and three A-T) according to the predicted Mfold structure (Figure 4). Only thymine and guanine imino protons involved in this hydrogen bonding are expected to be visible in the ^1^H NMR spectrum (Figure 5). The three peaks observed between 13.2 and 13.8 ppm (Figure 6A) correspond closely in chemical shift to those observed previously for thymine imino protons (49) and could therefore represent the three thymine imino protons involved in Watson-Crick hydrogen bindings in Aptamer-2-40mer (45). Furthermore, NOE correlations were observed between resonances 13.37 and 7.30 ppm, 13.49 and 7.82 ppm, and 13.54 and 7.26 ppm (Figure 6B). These are proposed to correspond to through-space interactions between the thymine N3**H** and the adenine C2**H** which, in Watson-Crick base pairing, is known to be significant (44,50). Therefore, the ^1^H NMR and NOE results are consistent with the three A-T base pairs proposed by Mfold to be present in the Aptamer-2-40mer stem in the favored structural form (Figure 4). Due to the similar chemical environment of thymines T2 and T4, their imino chemical shifts are expected to be very similar. By deduction, the two imino protons observed at 13.49 and 13.54 ppm are tentatively assigned as those of thymines T2 and T4, and, at 13.37 ppm, the imino proton of thymine T8.

**Figure 6.**
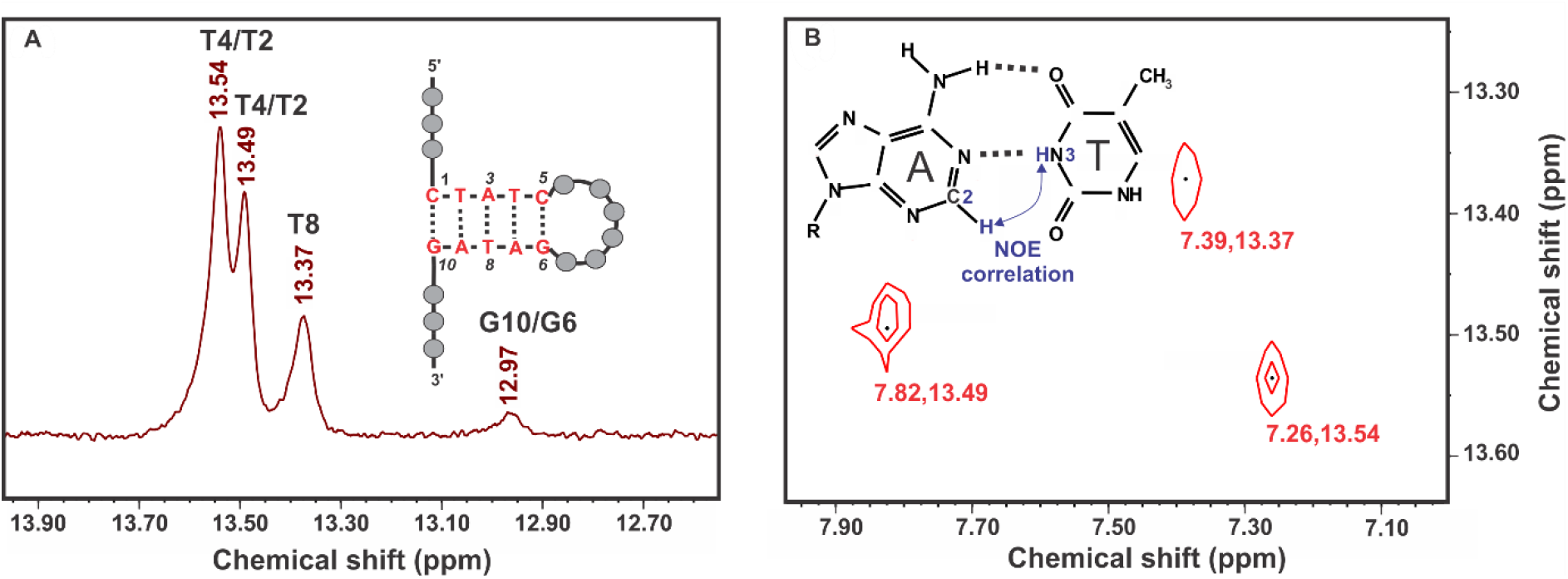
(A) Expansion of ^1^H NMR spectrum of Aptamer-2-40mer showing three thymine imino protons peaks (B) 2D NOESY correlations between the three imino protons and the three adenine C2 protons showing A-T base pairing.

The two G-C pairs in the structure predicted by Mfold are not clearly visible in the ^1^H NMR spectrum. This may be due to their position at the ends of the stem, meaning they are more prone to solvent exchange than the three NH protons of the thymines positioned in the middle of the stem. A weak resonance observed at 12.97 ppm may represent one imino NH from a guanine in the stem. This cannot be confirmed by NOE correlations in G-C base pairs due to the lack of a formamidine proton (or other similarly sited proton) in cytosine (44).The other chemical groups involved in hydrogen bonding are the amino groups of adenine, guanine, and cytosine (Figure 5), and these are observed in the 7.5-8.5 ppm region (Figure S5) (44,45).

This NMR-based structural assignment partially confirms the secondary structure predicted by Mfold in that the stem composition of three adenine-thymine Watson-Crick base pairs correspond to the spectral observations.

### ^1^H NMR spectroscopy reveals new imino proton signals upon Meth binding to Aptamer-2-40mer

^1^H NMR spectroscopy was used to investigate the binding mechanism and elucidate which part of Aptamer-2-40mer is involved in target recognition.

A comparison of the ^1^H NMR spectra of Aptamer-2-40mer in the absence and presence of Meth (saturated, 1 molar equivalent) shows the formation of four new imino resonances (peaks 5, 6, 7 and 8 in Figure 7) in the chemical shift region of hydrogen bonding for non-Watson-Crick base pairs (45). This is in line with studies by Neves et al. (49) and Jenison et al. (1) in which ligand addition caused additional resonances to appear, indicating that binding of Meth either induces a conformational change in the aptamer or stabilizes a Meth-bound aptamer conformation that otherwise exists in dynamic equilibrium with other unbound aptamer conformations.

**Figure 7.**
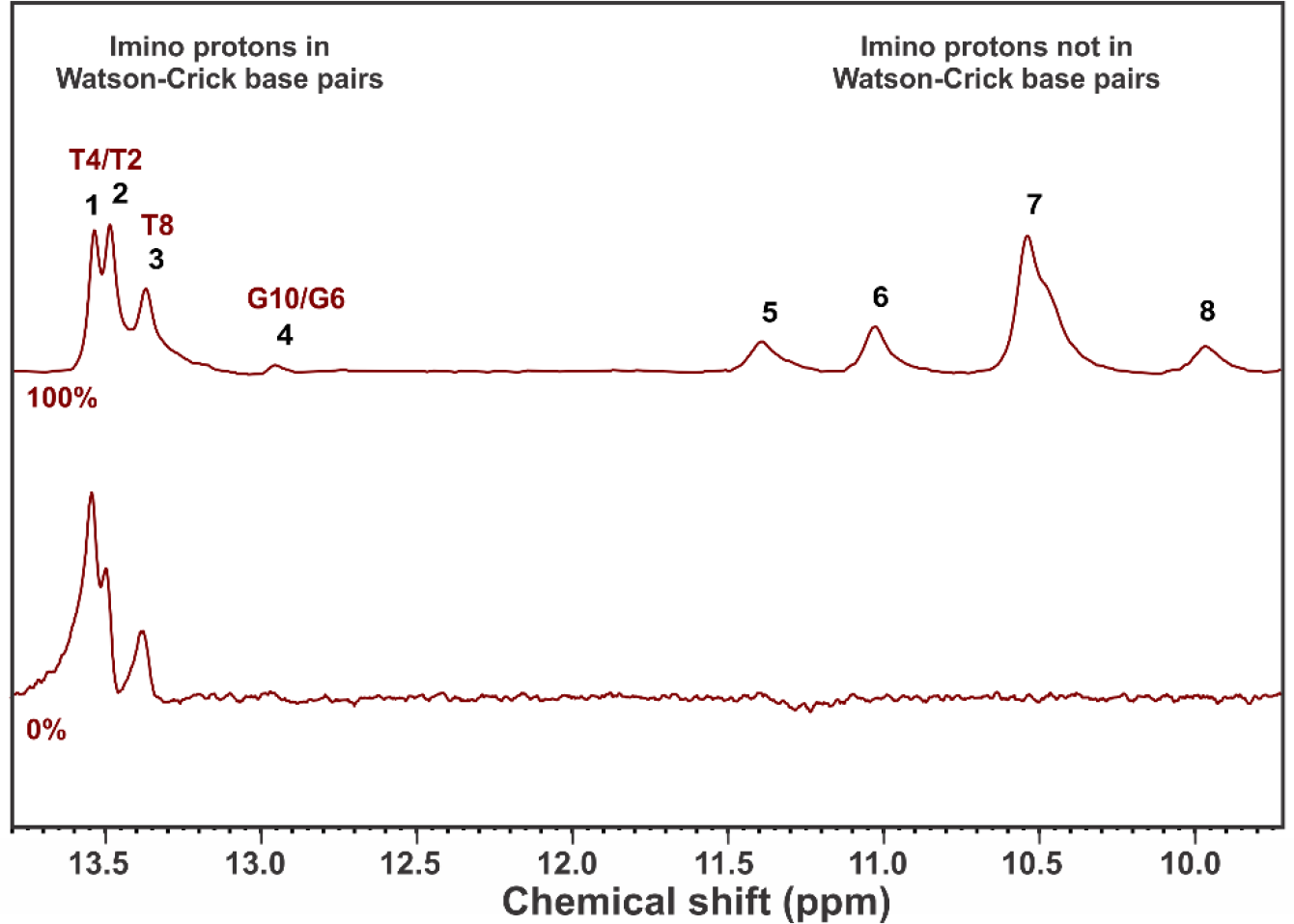
^1^H NMR spectra of Aptamer 2-40mer with molar loadings of Meth 0 % and 100 %.

Upon addition of 1 molar equivalent Meth, there was an apparent reduction in the intensity of peak 1 (assigned to the thymine imino proton involved in A-T2 or A-T4) relative to the other A-T resonances. This could be interpreted as a weakening of one of the two A-T interactions in the stem caused by Meth addition.

The appearance of peaks 5, 6, 7, and 8 (Figure 7), residing in the resonance region of imino protons not involved in Watson-Crick base pairs, leads to several hypotheses (*vide infra)*. Isothermal titration calorimetry was used, as described in the next section, to provide thermodynamics parameter to enable a better understanding about the creation of these new peaks and to propose a binding model related to the Meth-Aptamer-2-40mer complex formation.

### Isothermal titration calorimetry (ITC) confirms the importance of the stem and entropically driven binding

ITC was used for collecting thermodynamic properties and binding affinities related to Aptamer-2-40mer - Meth binding. Comparison of the original Aptamer-2-40mer with a mutated Aptamer-2-40mer sequence, where the base-paired stem is not able to form, highlights the importance of the stem and the resulting loop for binding. As shown in Figure 8A, the reaction is endothermic (heat consumption upon target addition) for the Aptamer-2-40mer, whereas the mutant lacking the ability to form a stem shows no variation of the calorimetric signal upon target addition in Figure 8B.

**Figure 8.**
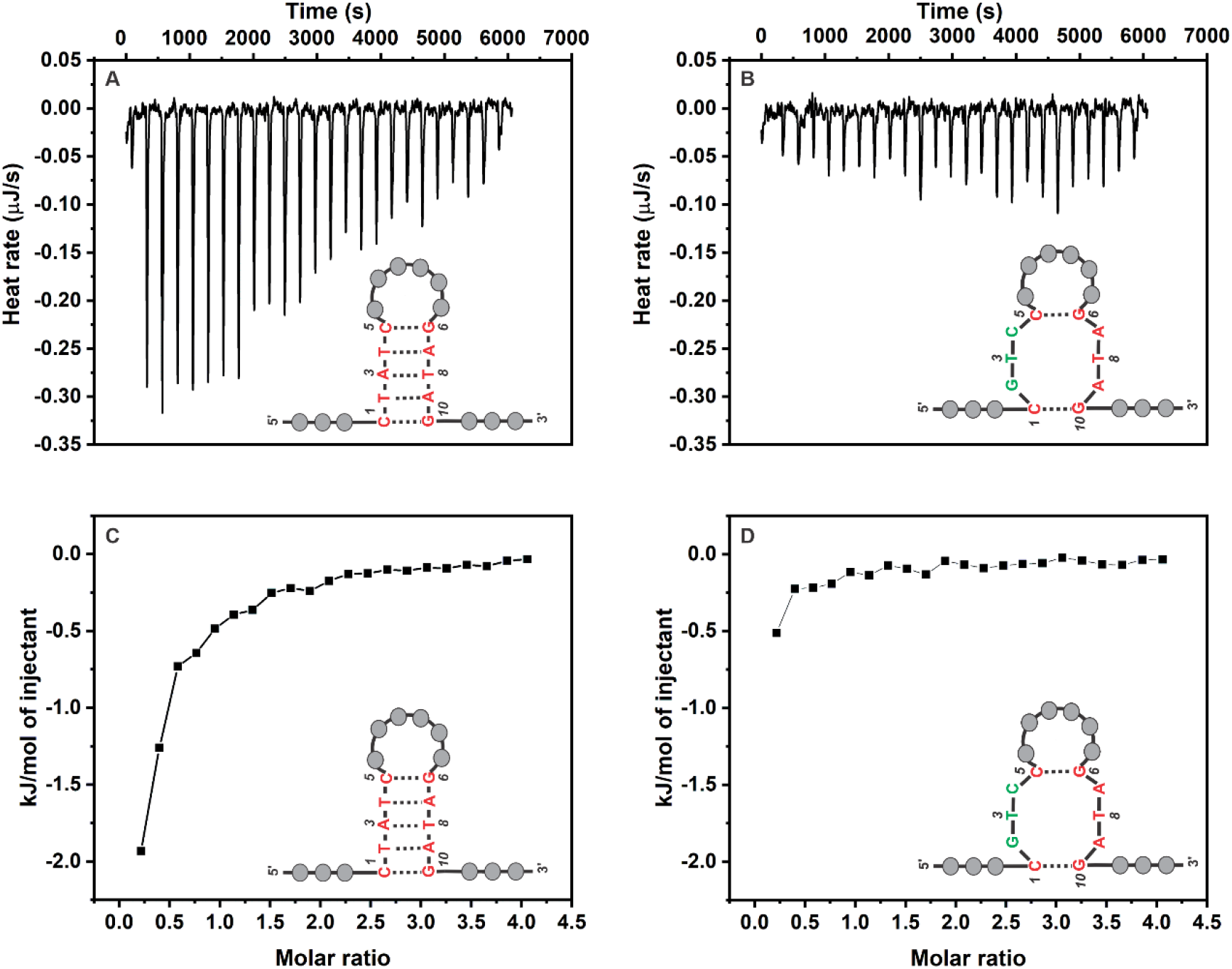
ITC results for the Aptamer-2-40mer and the mutated Aptamer-2-40mer. The heat profiles for (A) Aptamer-2-40mer and (B) mutated Aptamer-2-40mer are represented. The integrated heat profiles are represented for (C) Aptamer-2-40mer and (D) mutated Aptamer-2-40mer. The heat profiles presented have been corrected for the heat of dilution of the titrant and for the heat of dilution of the aptamer.

The mutated Aptamer-2-40mer was unable to bind with Meth (Figure 8B and 8D), which was attributable to the fact that the two remaining Watson-Crick base pairs cannot stabilize the stem formed in Aptamer-2-40mer. This experiment shows that disrupting the stem led to a full loss of binding, allowing us to conclude the stem is important to the specific binding event.

The enthalpic signal is weak and slightly endothermic (unfavorable contribution to the overall free energy of the aptamer-Meth complex), indicating disruption of non-covalent bonds (hydrogen bonds between the surrounding water and the aptamer surface) during the interaction between the ligand and the aptamer (13). Insufficient new hydrogen bonding interactions are created to overcome the enthalpy penalty. The weak calorimetric signal makes it hard to reliably extract *K_D_* and *ΔS* but in the absence of strong favorable enthalpic term, binding between Aptamer-2-40mer and Meth must be entropically driven.

The entropic component makes a favorable contribution to the overall free energy of the aptamer-Meth complex. The dominant contributors to the entropic effect could be the release of ordered solvated water molecules and counter-ions during ligand binding (51,52). A large favorable entropy component has previously been observed in hydrophobic binding interactions (13,53–55). Indeed, when ligand binding is associated with a large water gain entropy, the energy decrease (related to the attractive interactions between the aptamer and the Meth) is canceled out by the energy increase originating from the energetic dehydration effect upon binding (51). Given the complementary charges of the cationic Meth ligand and the anionic phosphate backbone of Aptamer-2-40mer, we can suggest that an electrostatic attraction may help to capture Meth molecules and position them near the stem region where hydrophobic interactions can be manifest. A related thermodynamic model (unfavorable binding enthalpy with a favorable binding entropy) has been previously proposed for the adsorption of the anionic surface of cytochrome *b_5_* on a positive charged anion-exchange chromatography (56). In both cases, an unfavorable overall binding enthalpy is compensated by a strong entropic contribution, and electrostatic interactions influence the nature of binding.

### Combination of spectroscopic and calorimetric data to establish a conformational selection binding model

By linking the results obtained from CD, NMR and ITC measurements, a binding model can be used to explain the Aptamer-2-40mer interaction with Meth. The three main binding models related to a bio-receptor – ligand interaction were displayed in Figure 1.

From the ^1^H NMR experiment, the creation of four imino proton peaks not involved in Watson-Crick base pairs upon Meth addition was observed (peaks 5, 6, 7 and 8 in Figure 7). The formation of the new peaks in the ^1^H NMR spectrum could result from intra-aptamer H-bonding (consistent with IF or CS binding models) or from other imino protons resonances such as H-bond of Meth-Aptamer, N**H** of Aptamer only or N**H_2_** of Meth only (consistent with IF, CS and LAK binding model). Two scenarios for the formation of these peaks can be considered:

[1] Meth is a hydrogen bond donor (Figure 2) and, thus, one H-bond may be formed between Meth and an oxygen or a nitrogen atom of a nucleotide base. This H-bond may have a chemical shift in the region of 10–11 ppm, which is the region of the imino protons not involved in Watson-Crick base pairs. In the case of a theophylline aptamer, Zimmermann et al. (57) observed the formation of a pseudo-base pair between the cytosine and the purine-like theophylline in the appearance of an H-bonding resonance in the region of 15 ppm. The Meth H-bonding interaction with an aptamer nucleotide will resonate in a more upfield region due to the lower electronegative environment provided by the Meth molecule. In addition, one of the Watson-Crick base pairs from the stem might be disrupted upon Meth addition and, therefore, one of the imino protons remains free in solution. It is still structurally protected by the stem from solvent exchange effects; thus, it is visible in the ^1^H NMR spectrum and resonates in the non-Watson-Crick region. Two signals are expected to appear in the 10.0–11.5 ppm region.

[2] No H-bond is formed between the Meth and the aptamer; however, the amine group can be resolved because Meth may interact with the aptamer in a specific cavity where protonated amine group of the Meth is protected from solvent exchange. The cationic amine group of Meth has two hydrogens that are not visible in the ^1^H NMR spectrum of the unbound ligand due to proton exchange with the aqueous environment but may become visible in the bound form (Figure S4). Neves et al. (49) showed that, when bound with its aptamer, the cocaine imino proton became visible in the 10 ppm region of the^1^H NMR spectrum. A similar effect may also occur with the Meth amino protons. Indeed, if Meth interacts with a specific cavity of the aptamer, proton exchange with the aqueous environment would similarly decrease due to the structural protection of the aptamer, and possibly resolve the two amino proton resonances. The electron demand is similar for Meth and cocaine, nevertheless, as Meth is a secondary amine and cocaine a tertiary amine the chemical shift will differ slightly. One of the Watson-Crick base pairs from the stem may be disrupted and, therefore, one of the imino protons is free in solution (visible as explained in the first scenario) and is resonated in this region. Three signals are expected to appear in the 10.0–11.5 ppm region.

Neither of these scenarios explain the four new NMR peaks observed. Consequently, some of the four peaks observed may be related to intra-aptamer H bonds and, thus, it indicates that different structures of the aptamer are present in solution before and after target addition. Consequently, the LAK binding model is ruled out. The structural observations obtained with NMR are in line with the CS and IF binding models.

A main difference between the CS and the IF binding models is the number of conformational states present initially in solution (6,13–15). The IF binding model assumes an initial non-binding conformation present in solution; by adding the target to a 1:1 molar ratio (aptamer:target), it is considered that a new binding state is present and no non-binding conformation remains in solution. From a thermodynamic perspective, the IF binding model would require a strong attractive interaction (i.e. enthalpic) to drive this structural change (13,58,59). This is inconsistent with the lack of enthalpic attraction observed in the ITC experiment so we can suppose that the IF binding model for the Aptamer-2-40mer - Meth interaction can be ruled out.

Moreover, the CD experiments support this suggestion because no significant change in the duplex structure upon ligand addition is observed (Figure 3). Furthermore, in the ^1^H NMR experiment, we observed that the three peaks (1, 2 and 3 in Figure 6) related to the stem are present before the target addition. Despite the modest decrease in intensity of peak 1, all are still present after the target addition. No new resonance peaks related to Watson-Crick base pair formation are visible after Meth addition. These two observations provide additional evidence that no major aptamer structural changes occur upon Meth addition.

Finally, the CS (conformational-selection) binding model is considered. Different conformations of the aptamer exist in a dynamic equilibrium in solution as shown in Figure 1B (13,14,60). Meth binds specifically to one of these conformations. At 0% Meth, the aptamer conformations are in a dynamic equilibrium where *k′_rev_*>*k′_for_* (unbound conformation is favored and thus is observed at 0% Meth). The addition of Meth causes the binding state to be favored in solution *k′_on_*>*k′_rev_*; therefore, the ^1^H NMR spectrum upon addition of one equivalent (saturated) represents the binding state with additional non-Watson-Crick resonances. Because the loop region contains enough guanine or thymine residues to form non-Watson-Crick base pairs, it is hypothesized that these interactions are situated in the loop region. ITC experiments support this hypothesis as no signal is measured for the mutated Aptamer-2-40mer where the loop is not formed. Furthermore, in support of the thermodynamic data collected, a mixture of binding and non-binding state conformation is present in solution. By adding the target, the equilibrium between the two states shifts to the binding state conformation. Comparatively to the IF binding mode, the equilibrium shift is less significant due to the initial presence of binding state aptamer. Thermodynamically, this equilibrium switch may not afford a strong attractive interaction (enthalpic in contrast to the IF binding mode) and, thus, is consistent with an entropically driven binding event.

### Molecular modeling supports the proposed binding structure

Following the experimental determination of Meth binding to the Aptamer-2-40mer, molecular docking was used to rationalize the proposed binding hypotheses.

First, a 3D model of the Aptamer-2-40mer was constructed using RNAcomposer (28). Mfold (26) and RNAstructure (27) provided a 2D structure corresponding to experimental observations (26,27). A stem composed of the three A-T base pairs is predicted and conforms with NMR experimental evidence. Additionally, with the RNAstructure software, the partition function calculation (61) predicted base pair probabilities. By using the partition function for the Aptamer-2-40mer, it appears that the five base pairs predicted by Mfold and represented in the Aptamer-2-40mer structure (Figure 4) have a high probability of formation (Figure S5). Finally, the 2D structure and DNA sequence have been used to generate the 3D structure with RNAcomposer.

Exhaustive docking of Meth revealed several binding poses homogeneously distributed within the major groove of the stem region (Figure 9A). All poses predict the placement of the cationic amine group of Meth near C5, T4, A′ (a nucleotide within the loop), T′′ (a nucleotide within the loop), T′′′ (a nucleotide within the loop), G6, and the aromatic ring near T2, A3, and A7. Conformational clustering of the 100 poses gave two distinct clusters: the highest scored was least populated, with 12 poses (**C1**), whilst the most populated (**C2**) contained 88 poses that scored slightly worse (see Figure 9B for score distributions). Analysis of **C1** (Figure 9C) shows the cationic amine of Meth forming a salt bridge with the T′′ phosphate backbone. The remainder of the Meth framework (from methyl-9 to the aromatic ring) resides along a column of polar moieties from the surrounding bases (G6 carbonyl, C5 amine, T4 carbonyl, A7 amine, A3 amine, T8 carbonyl). For **C2** (Figure 9D), the cationic amine retains the ionic bond with the T′′ phosphate backbone, but the methyl-9 of Meth is positioned towards the T′′′ thymine methyl group rather than the T4 thymine methyl as seen in **C1**. Overall, the docking study suggests Meth could favorably occupy the pocket by forming ionic and hydrophobic interactions with the surrounding environment. These findings are in line with those experimentally deduced, and both highlight the importance of the stem region for Meth binding. The predicted binding modes may also support the hypothesis that the magnitude change of peak 1 in the ^1^H NMR spectrum upon Meth treatment (in Figure 7) corresponds to partial disruption of H-bonding between A-T2 or A-T4 base pairs. Furthermore, the hydrophobic and ionic interactions between the phosphate backbone and Meth correlate with the thermodynamic parameters obtained by ITC (Figure 8).

**Figure 9.**
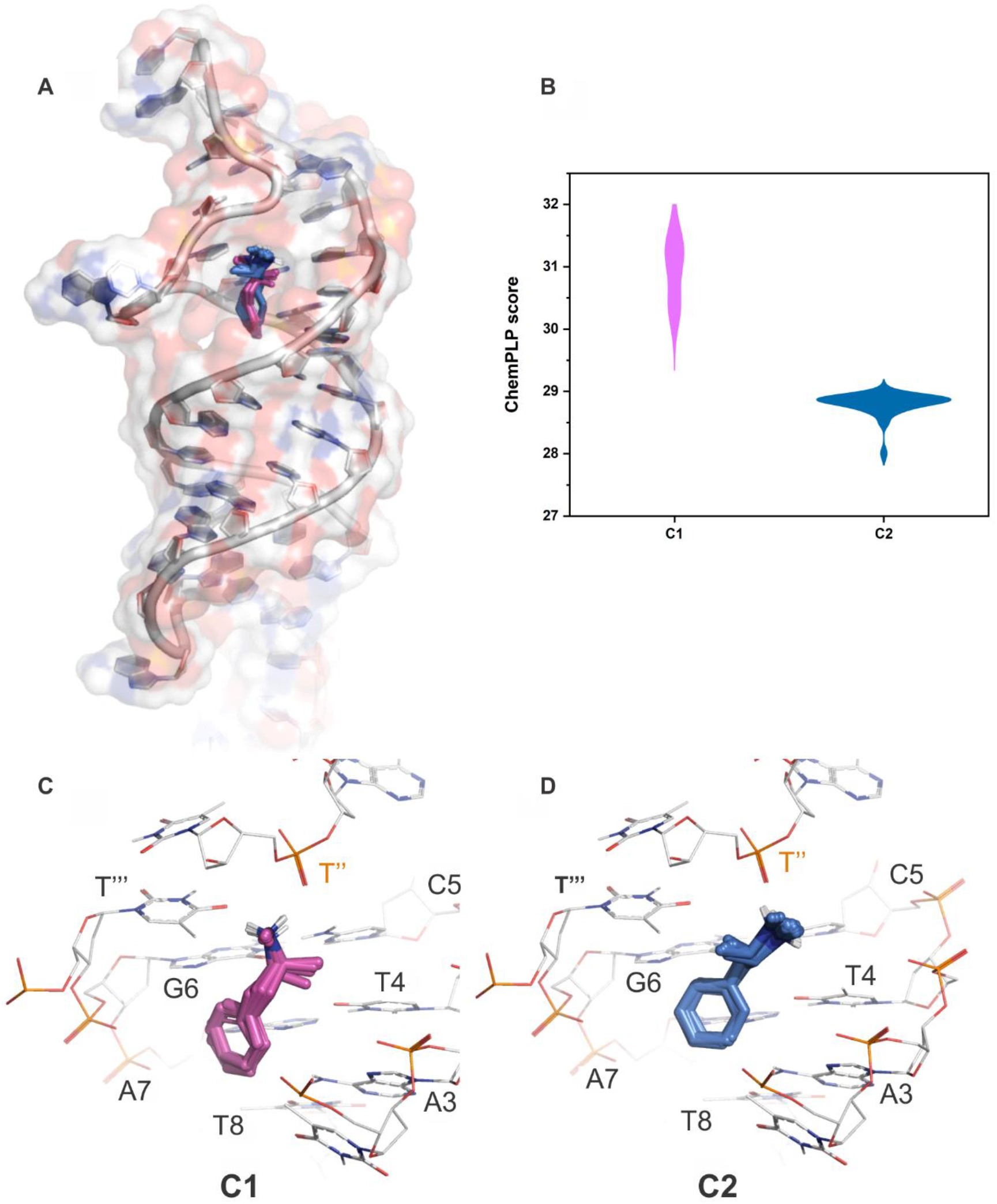
Molecular docking of Meth with Aptamer-2-40mer. (A) All 100 docking poses of Meth in the hydrophobic pocket are shown in stick form, the aptamer is shown in cartoon form with a transparent surface rendered. (B) A violin plot of the docking scores of each cluster. (C) The 12 poses of cluster_1 **C1** (Meth in stick form) in the pocket with surrounding nucleotides shown in wire form. (D) The 88 poses of cluster_2 **C2** (Meth in stick form) in the pocket with surrounding nucleotides shown in wire form.

## CONCLUSIONS

Through a combination of CD, NMR, and ITC experiments, a binding mechanism for the Meth – Aptamer-2-40mer complex is proposed and supported by molecular modeling simulations. Specifically, no significant structural change has been detected with CD experiments upon addition of Meth. NMR experiments indicate a change in the imino region for Aptamer-2-40mer in solution upon Meth addition that is proposed to consist of a form where new non-Watson-Crick base pairs are present in the loop (potentially arising through a conformational-selection binding process). Moreover, a partial weakening of one of the A-T2 or A-T4 Watson-Crick pairs is tentatively interpreted from the NMR spectra to occur upon target addition. ITC experiments demonstrate the importance of the stem for the binding event. Indeed, by removing the stem, no binding event occurs. Thermodynamic parameters collected with ITC provided information suggesting an entropically driven binding event. Molecular modeling supports the proposed hypotheses in the preferred positions of Meth calculated to be within the major groove of the stem and with interactions to the phosphate backbone of the loop.

This approach is suited to study of other aptamer-small molecule complexes as a characterization protocol prior to development of an aptasensor platform. Following the findings related to the binding region localization as well as the structural switch magnitude of the aptamer upon the target addition, the best transduction strategy needs to be considered. The small magnitude of the conformational switch and accompanying enthalpic change indicate that Affinity Matrix SELEX may not provide the optimal aptamer structures for detection of monofunctional small molecule targets. Other SELEX protocols must be investigated for this purpose.

## Supporting information

Supplemental equation, table and figures

## DATA AVAILABILITY

All aptamer sequences used in the study are reported in the SI.

## SUPPLEMENTARY DATA

Supplementary Data are available online.

## REFERENCES

1. Jenison, R.D., Gill, S.C., Pardi, A. and Polisky, B. (1994) High-resolution molecular discrimination by RNA. Science, 263, 1425.

2. Tuerk, C. and Gold, L. (1990) Systematic Evolution of Ligands by Exponential Enrichment: RNA Ligands to Bacteriophage T4 DNA Polymerase. Science, 249, 505–510.

3. Ellington, A.D. and Szostak, J.W. (1990) In vitro selection of RNA molecules that bind specific ligands. Nature, 346, 818.

4. Jayasena, S.D. (1999) Aptamers: An Emerging Class of Molecules That Rival Antibodies in Diagnostics. Clinical Chemistry, 45, 1628–1650.

5. Zhou, J. and Rossi, J. (2017) Aptamers as targeted therapeutics: current potential and challenges. Nat Rev Drug Discov, 16, 181–202.

6. Munzar, J.D., Ng, A. and Juncker, D. (2018) Comprehensive profiling of the ligand binding landscapes of duplexed aptamer families reveals widespread induced fit. Nature Communications, 9, 343.

7. Feagin, T.A., Maganzini, N. and Soh, H.T. (2018) Strategies for Creating Structure-Switching Aptamers. ACS Sens, 3, 1611–1615.

8. Zhang, Z., Oni, O. and Liu, J. (2017) New insights into a classic aptamer: binding sites, cooperativity and more sensitive adenosine detection. Nucleic Acids Res, 45, 7593–7601.

9. Wilson, B.D., Hariri, A.A., Thompson, I.A.P., Eisenstein, M. and Soh, H.T. (2019) Independent control of the thermodynamic and kinetic properties of aptamer switches. Nature Communications, 10, 5079.

10. Xiao, Y., Lubin, A.A., Heeger, A.J. and Plaxco, K.W. (2005) Label-Free Electronic Detection of Thrombin in Blood Serum by Using an Aptamer-Based Sensor. Angewandte Chemie, 117, 5592–5595.

11. Stojanovic, M.N., de Prada, P. and Landry, D.W. (2001) Aptamer-Based Folding Fluorescent Sensor for Cocaine. Journal of the American Chemical Society, 123, 4928–4931.

12. Baker, B.R., Lai, R.Y., Wood, M.S., Doctor, E.H., Heeger, A.J. and Plaxco, K.W. (2006) An Electronic, Aptamer-Based Small-Molecule Sensor for the Rapid, Label-Free Detection of Cocaine in Adulterated Samples and Biological Fluids. Journal of the American Chemical Society, 128, 3138–3139.

13. Du, X., Li, Y., Xia, Y.-L., Ai, S.-M., Liang, J., Sang, P., Ji, X.-L. and Liu, S.-Q. (2016) Insights into Protein-Ligand Interactions: Mechanisms, Models, and Methods. International journal of molecular sciences, 17, 144.

14. Ma, B., Kumar, S., Tsai, C.J. and Nussinov, R. (1999) Folding funnels and binding mechanisms. Protein Eng, 12, 713–720.

15. Csermely, P., Palotai, R. and Nussinov, R. (2010) Induced fit, conformational selection and independent dynamic segments: an extended view of binding events. Trends Biochem Sci, 35, 539–546.

16. Fischer, E. (1894) Einfluss der Configuration auf die Wirkung der Enzyme. Berichte der deutschen chemischen Gesellschaft, 27, 2985–2993.

17. Hermann, T. and Patel, D.J. (2000) Adaptive Recognition by Nucleic Acid Aptamers. Science, 287, 820–825.

18. Koshland, D.E. (1958) Application of a Theory of Enzyme Specificity to Protein Synthesis. Proc Natl Acad Sci U S A, 44, 98–104.

19. Monod, J., Wyman, J. and Changeux, J.-P. (1965) On the nature of allosteric transitions: a plausible model. J Mol Biol, 12, 88–118.

20. Gluckman, P. (2018) In Advisor, O. o. t. P. M. s. C. S. (ed.).

21. Li, S., Clarkson, M. and McNatty, K. (2020) Selection and characterisation of triclosan-specific aptamers using a fluorescence microscope-imaging assay. Analytical and Bioanalytical Chemistry, 412, 7285–7294.

22. Wishart, D.S., Bigam, C.G., Yao, J., Abildgaard, F., Dyson, H.J., Oldfield, E., Markley, J.L. and Sykes, B.D. (1995) 1H, 13C and 15N chemical shift referencing in biomolecular NMR. Journal of Biomolecular NMR, 6, 135–140.

23. Jeener, J., Meier, B.H., Bachmann, P. and Ernst, R.R. (1979) Investigation of exchange processes by two – dimensional NMR spectroscopy. The Journal of Chemical Physics, 71, 4546–4553.

24. Piotto, M., Saudek, V. and Sklenář, V. (1992) Gradient-tailored excitation for single-quantum NMR spectroscopy of aqueous solutions. Journal of Biomolecular NMR, 2, 661–665.

25. Willcott, M.R. (2009) MestRe Nova. Journal of the American Chemical Society, 131, 13180–13180.

26. Zuker, M. (2003) Mfold web server for nucleic acid folding and hybridization prediction. Nucleic Acids Research, 31, 3406–3415.

27. Reuter, J.S. and Mathews, D.H. (2010) RNAstructure: software for RNA secondary structure prediction and analysis. BMC Bioinformatics, 11, 129.

28. Popenda, M., Szachniuk, M., Antczak, M., Purzycka, K.J., Lukasiak, P., Bartol, N., Blazewicz, J. and Adamiak, R.W. (2012) Automated 3D structure composition for large RNAs. Nucleic Acids Research, 40, e112–e112.

29. Schrödinger. (2020) In LLC (ed.), New York, NY.

30. Jones, G., Willett, P., Glen, R.C., Leach, A.R. and Taylor, R. (1997) Development and validation of a genetic algorithm for flexible docking. J Mol Biol, 267, 727–748.

31. Korb, O., Stützle, T. and Exner, T.E. (2009) Empirical Scoring Functions for Advanced Protein-Ligand Docking with PLANTS. Journal of Chemical Information and Modeling, 49, 84–96.

32. Schrödinger. (2019) In LLC (ed.), New York, NY.

33. Kelley, L.A., Gardner, S.P. and Sutcliffe, M.J. (1996) An automated approach for clustering an ensemble of NMR-derived protein structures into conformationally related subfamilies. Protein Engineering, Design and Selection, 9, 1063–1065.

34. Cai, S., Yan, J., Xiong, H., Liu, Y., Peng, D. and Liu, Z. (2018) Investigations on the interface of nucleic acid aptamers and binding targets. Analyst, 143, 5317–5338.

35. Zipper, H., Brunner, H., Bernhagen, J. and Vitzthum, F. (2004) Investigations on DNA intercalation and surface binding by SYBR Green I, its structure determination and methodological implications. Nucleic Acids Res, 32, e103.

36. Kong, L., Xu, J., Xu, Y., Xiang, Y., Yuan, R. and Chai, Y. (2013) A universal and label-free aptasensor for fluorescent detection of ATP and thrombin based on SYBR Green I dye. Biosens Bioelectron, 42, 193–197.

37. Watson, J.D. and Crick, F.H.C. (1953) Molecular Structure of Nucleic Acids: A Structure for Deoxyribose Nucleic Acid. Nature, 171, 737–738.

38. Vaughan, D.M.G.R.L.R.M.R. (1992), Vol. 211.

39. Bishop, G.R. and Chaires, J.B. (2002) Characterization of DNA Structures by Circular Dichroism. Current Protocols in Nucleic Acid Chemistry, 11, 7.11.11–17.11.18.

40. Gondeau, C., Maurizot, J.C. and Durand, M. (1998) Circular dichroism and UV melting studies on formation of an intramolecular triplex containing parallel T*A:T and G*G:C triplets: netropsin complexation with the triplex. Nucleic Acids Research, 26, 4996–5003.

41. Bing, T., Zheng, W., Zhang, X., Shen, L., Liu, X., Wang, F., Cui, J., Cao, Z. and Shangguan, D. (2017) Triplex-quadruplex structural scaffold: a new binding structure of aptamer. Sci Rep, 7, 15467.

42. Vorlickova, M., Kejnovska, I., Bednarova, K., Renciuk, D. and Kypr, J. (2012) Circular dichroism spectroscopy of DNA: from duplexes to quadruplexes. Chirality, 24, 691–698.

43. Li, H.-H., Wen, C.-Y., Hong, C.-Y. and Lai, J.-C. (2017) Evaluation of aptamer specificity with or without primers using clinical samples for C-reactive protein by magnetic-assisted rapid aptamer selection. RSC Advances, 7, 42856–42865.

44. Fürtig, B., Richter, C., Wöhnert, J. and Schwalbe, H. (2003) NMR Spectroscopy of RNA. ChemBioChem, 4, 936–962.

45. Wijmenga, S.S. and van Buuren, B.N.M. (1998) The use of NMR methods for conformational studies of nucleic acids. Progress in Nuclear Magnetic Resonance Spectroscopy, 32, 287–387.

46. Mirau, P.A., Smith, J.E., Chávez, J.L., Hagen, J.A., Kelley-Loughnane, N. and Naik, R. (2018) Structured DNA Aptamer Interactions with Gold Nanoparticles. Langmuir, 34, 2139–2146.

47. Leontis, N.B., Stombaugh, J. and Westhof, E. (2002) The non-Watson-Crick base pairs and their associated isostericity matrices. Nucleic acids research, 30, 3497–3531.

48. Hermann, T. and Westhof, E. (1999) Non-Watson-Crick base pairs in RNA-protein recognition. Chemistry & Biology, 6, R335–R343.

49. Neves, M.A.D., Slavkovic, S., Churcher, Z.R. and Johnson, P.E. (2017) Salt-mediated two-site ligand binding by the cocaine-binding aptamer. Nucleic acids research, 45, 1041–1048.

50. Lin, C.H., Wang, W., Jones, R.A. and Patel, D.J. (1998) Formation of an amino-acid-binding pocket through adaptive zippering-up of a large DNA hairpin loop. Chemistry & Biology, 5, 555–572.

51. Hayashi, T., Oshima, H., Mashima, T., Nagata, T., Katahira, M. and Kinoshita, M. (2014) Binding of an RNA aptamer and a partial peptide of a prion protein: crucial importance of water entropy in molecular recognition. Nucleic Acids Research, 42, 6861–6875.

52. Potty, A.S.R., Kourentzi, K., Fang, H., Schuck, P. and Willson, R.C. (2011) Biophysical characterization of DNA and RNA aptamer interactions with hen egg lysozyme. International Journal of Biological Macromolecules, 48, 392–397.

53. Sakamoto, T., Ennifar, E. and Nakamura, Y. (2018) Thermodynamic study of aptamers binding to their target proteins. Biochimie, 145, 91–97.

54. Fisher, H.F. and Singh, N. (1995), Methods in Enzymology. Academic Press, Vol. 259, pp. 194–221.

55. Archer, W.R. and Schulz, M.D. (2020) Isothermal titration calorimetry: practical approaches and current applications in soft matter. Soft Matter, 16, 8760–8774.

56. Gill, D.S., Roush, D.J. and Willson, R.C. (1994) Presence of a preferred anion-exchange binding site on cytochrome b5: structural and thermodynamic considerations. Journal of Chromatography A, 684, 55–63.

57. Zimmermann, G.R., Wick, C.L., Shields, T.P., Jenison, R.D. and Pardi, A. (2000) Molecular interactions and metal binding in the theophylline-binding core of an RNA aptamer. RNA, 6, 659–667.

58. Li, H., Xie, Y., Liu, C. and Liu, S. (2014) Physicochemical bases for protein folding, dynamics, and protein-ligand binding. Science China Life Sciences, 57, 287–302.

59. Chang, C.-E. and Gilson, M.K. (2004) Free Energy, Entropy, and Induced Fit in Host-Guest Recognition: Calculations with the Second-Generation Mining Minima Algorithm. Journal of the American Chemical Society, 126, 13156–13164.

60. Paul, F. and Weikl, T.R. (2016) How to Distinguish Conformational Selection and Induced Fit Based on Chemical Relaxation Rates. PLOS Computational Biology, 12, e1005067.

61. Mathews, D.H. (2004) Using an RNA secondary structure partition function to determine confidence in base pairs predicted by free energy minimization. RNA, 10, 1178–1190.

