## Supplemental equation, table and figures for "Unravelling the binding mode of a methamphetamine aptamer: a spectroscopic and calorimetric investigation"

Binding constant equation related to the "**Lock and Key**" equilibrium is the following: $K_D{}^{obs} = \frac{k_{off}}{k_{on}}$. Binding constant equations related to the "**Conformational Selection**" equilibrium are the following: $K_D{}^{obs} = K_{disso}\,(1 + K_{iso})$ where $K_{disso} = \frac{k'_{off}}{k'_{on}}$ and $K_{iso} = \frac{k'_{rev}}{k'_{for}}$. Binding constant equations related to the "**Induced Fit**" equilibrium are the following: $K_D{}^{obs} = K_{disso}\,\frac{K_{iso}}{(1+K_{iso})}$ where $K_{disso} = \frac{k''_{off}}{k''_{on}}$ and $K_{iso} = \frac{k''_{rev}}{k''_{for}}$.

**Equation S1.** Binding constant equations related to the Lock-and-Key, Conformational Selection, and Induced Fit binding models.

| Aptamer identification | Sequence 5’-3’ |
|---|---|
| **Aptamer-1** | ATACGAGCTTGTTCAATAGCGTTTAGGCGTTCATTCATCCCGCTATCTGGCTGTATCGTGATAGTAAGAGCAATC |
| **Aptamer-2** | ATACGAGCTTGTTCAATAGCGTTTCTATCTGGCTGTATCGTGATAGTAAGAGCACTAATGATAGTAAGAGCAATC |
| **Aptamer-3** | ATACGAGCTTGTTCAATAGCGTTTAGCGTTCAATTCATCCCGCTATCCTGGCTGTATCGTGATAGTAGAACAATC |
| **Aptamer-4** | ATACGAGCTTGTTCAATAGCGTTTTACGTTCAATTCATCCCGCTATCTGGCTGTATCGTGATAGTAAGAGCAATC |
| **Aptamer-2-40mer** | GCGTTTCTATCTGGCTGTATCGTGATAGTAAGAGCACTAA |
| **Mutated-Aptamer-2-40mer** | GCGTTTCGTCCTGGCTGTATCGTGATAGTAAGAGCACTAA |

**Table S1.** Full Meth-aptamer sequences used in the study. Buffer conditions are 2 mM Tris-HCl pH 7.5, 10 mM NaCl, 0.5 mM KCl, 0.2 mM $MgCl_2$ and 0.1 mM $CaCl_2$.

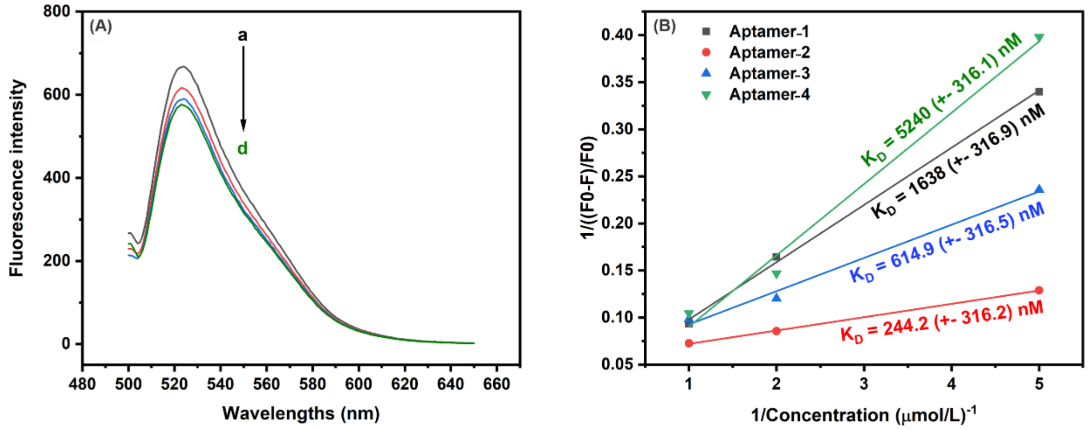

**Figure S1.** Meth titration (0 µM (a), 0.2 µM (b), 0.5 µM (c) and 1 µM (d)) for the Meth aptamers family. (A) A fluorescence decrease at 520 nm is monitored for Aptamer-2. (B) Linearization of the Langmuir isotherm for each aptamer.

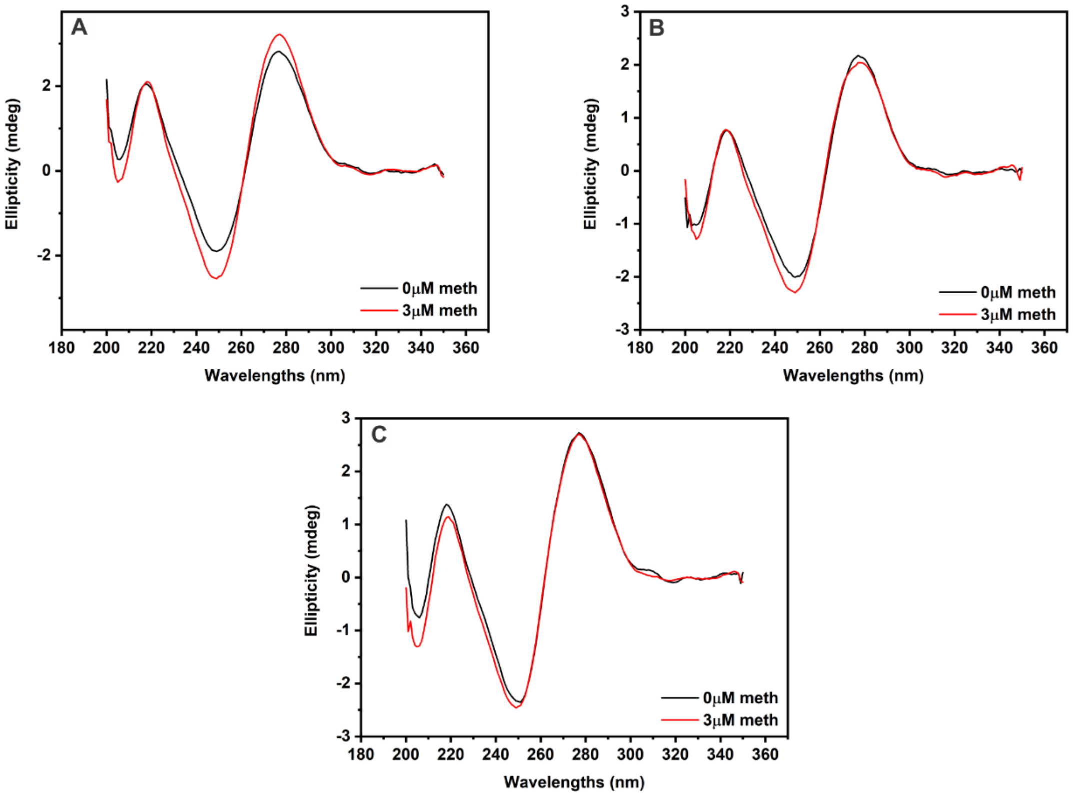

**Figure S2.** CD experiments for the three other Meth aptamers (A) Aptamer-1, (B) Aptamer-3 and (C) Aptamer-4.

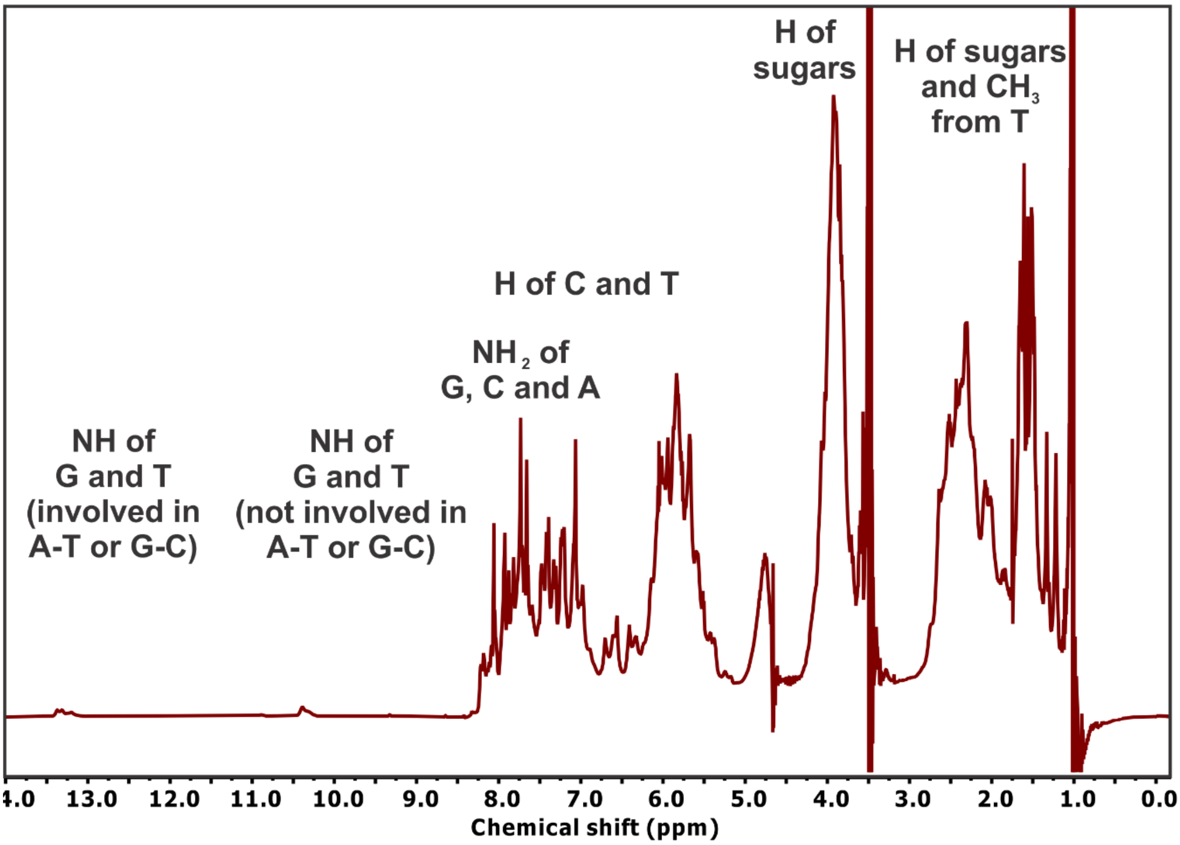

**Figure S3.** Aptamer-2-40mer $^1$H-NMR spectrum.

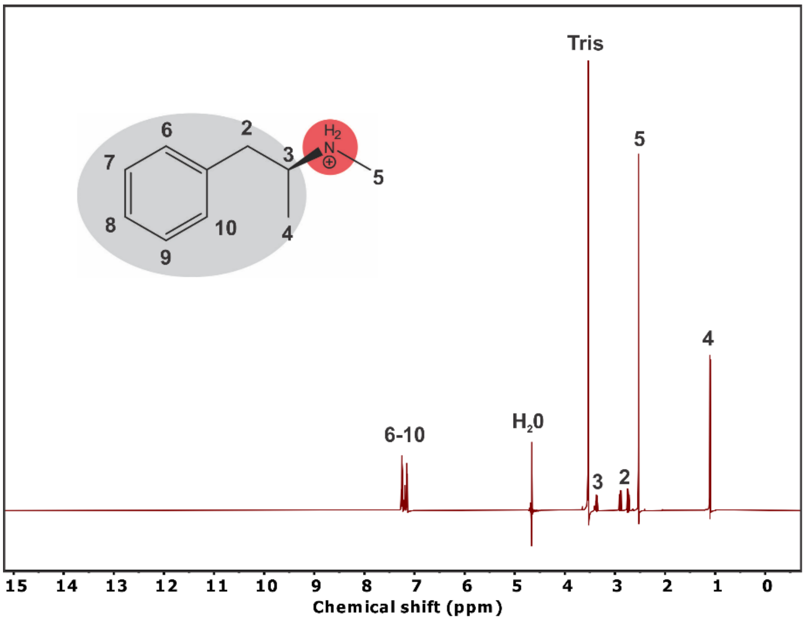

**Figure S4.** Meth $^1$H-NMR spectrum.

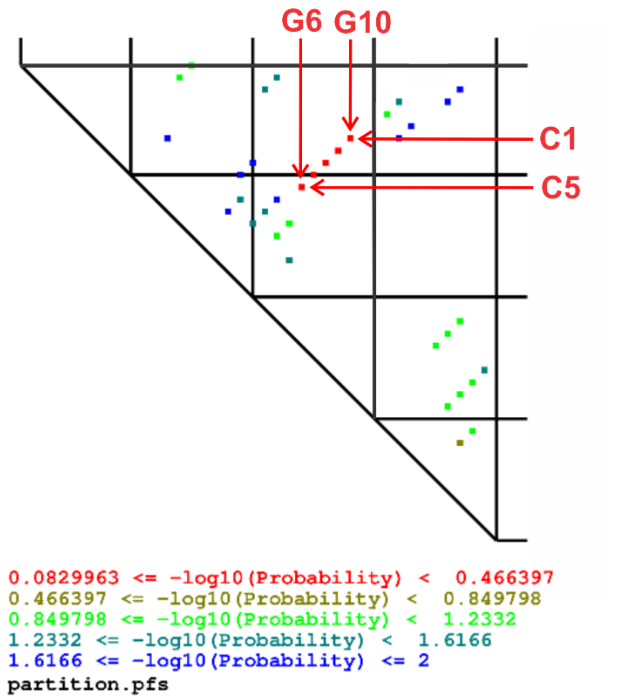

**Figure S5.** Partition results for Aptamer-2-40mer obtained from *RNAstructure*.
